# Genome-wide association between transcription factor expression and chromatin accessibility reveals chromatin state regulators

**DOI:** 10.1101/043414

**Authors:** David Lamparter, Daniel Marbach, Rico Rueedi, Sven Bergmann, Zoltán Kutalik

## Abstract

To better understand genome regulation, it is important to uncover the role of transcription factors in the process of chromatin structure establishment and maintenance. Here we present a data-driven approach to systematically characterize transcription factors that are relevant for this process. Our method uses a linear mixed modeling approach to combine data sets of transcription factor binding motif enrichments in open chromatin and gene expression across the same set of cell lines. Applying this approach to the ENCODE data set we confirm already known and imply numerous novel transcription factors in playing a role in the establishment or maintenance of open chromatin.

## Introduction

In higher eukaryotes, certain sequence-specific transcription factors (TFs), which we will call *chromatin state regulators* (CSRs), are responsible for establishing and maintaining open chromatin configurations (Zaret and Carroll 2011; Morris et al. 2013). CSRs therefore play a fundamental role in transcriptional regulation, because open chromatin configurations are necessary for additional TFs to bind and transcriptionally activate target genes.

CSRs that can bind closed chromatin and open up chromatin are called pioneer TFs (Iwafuchi-Doi and Zaret 2014). The comprehensive identification of pioneer TFs with high confidence still needs further research. While some pioneers are well studied, others have only preliminary evidence, or are only computationally predicted. Some well studied examples include *FOXA1*, whose winged helix domains disrupt DNA-histone contacts, and *POU5F1, SOX2* and *KLF4*, which are used in production of iPSC cell lines (Cirillo and Zaret 2007; Soufi et al. 2014). Further pioneer TFs such as *ASCL1, SPI1* and the *GATA* factors are used in transdifferentiation, and *PAX7* is involved in pituitary melanotrope development (Vierbuchen et al. 2010; Feng et al. 2008; Cirillo et al. 2002a; Budry et al. 2012). However, not all pioneer TFs are involved in development and cell conversions. *CLOCK/BMAL1* are part of the circadian clock and the tumor suppressor *TP53* is involved in the cell cycle, while its close homolog *TP63* is involved in skin development (Menet et al. 2014; Sammons et al. 2015; Koster 2010).

Recent studies suggest that maintaining open chromatin is a dynamic process with pioneer and other TFs binding and unbinding rapidly and continually recruiting additional chromatin remodelling factors that are not sequence specific (Morris et al. 2013; Voss and Hager 2014; Voss et al. 2011; Nagaich et al. 2004). TFs vary in their ability to recruit particular remodelling factors, for example the TFs *STAT5A/B* and *MYOG* motifs enrich in binding sites of the *SWI/SNF* remodelling complex but not in *ISWI* remodelling complex binding sites whereas *YY1* motifs were found exclusively *in ISWI* complex binding sites (Morris et al. 2013). A natural question then is which TFs are relevant to maintain open chromatin and can therefore be called CSRs.

One approach to test whether a given TF is a CSR is to perform a knock-down of this TF followed by an open chromatin assay to see whether chromatin regions containing the respective motif preferentially change from open to closed (Harrison et al. 2011). However, this approach is very time consuming because it requires a separate knock-down experiment for each TF. To define pioneer TFs specifically, one can check if the TF has the ability to bind nucleosomal DNA *in vitro* and validate the results *in vivo (Cirillo et al. 2002b*). Recently, a computational method called *Protein interaction Quantification (PIQ*) has been published that aims to recover pioneer TFs by estimating both TF binding and ensuing chromatin changes from the same Dnase1 hypersensitivity (DHS) experiments (Sherwood et al. 2014). However, *PIQ* did not predict some well known pioneer TFs such as *FOXA1, SOX2* and *POU5F1* showing that further improvements are possible (Iwafuchi-Doi and Zaret 2014).

Here we introduce a data driven approach to determine CSRs (Fig. 1). Our approach relies on the joint analysis of a large collection of DHS and coordinated gene expression data to estimate TF activity independently of DHS data. We first define the *motif accessibility score* for a given TF for each cell line based on the enrichment of its binding motif in regions with open chromatin. Then, we associate these scores with gene expression values across all available cell lines. We used our approach on data generated as part of the ENCODE project (Bernstein et al. 2012; Thurman et al. 2012). This uncovered numerous TFs whose motif accessibility is robustly associated with mRNA expression across 109 cell lines suggesting a role in the establishment or maintenance of open chromatin. Also, we see that our uncovered TFs are strongly enriched for known pioneer TFs. This suggests that the TFs we identified are good candidates for CSRs.

**Figure 1:**
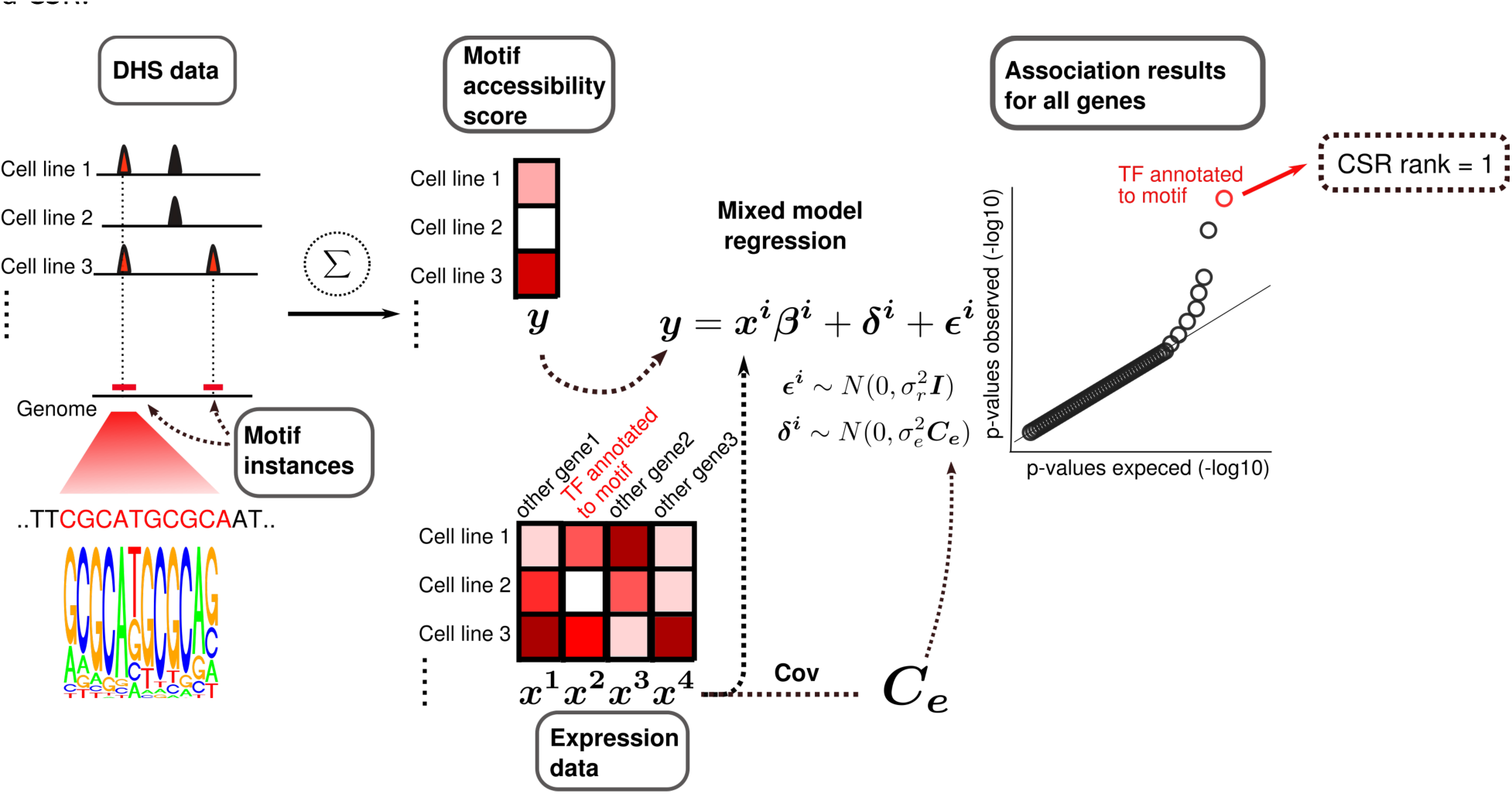
Mixed model approach for identification of chromatin state regulators. Overview of motif accessibility scoring and mixed modeling procedure. For a TF binding motif, we search for all its instances in the genome. For each cell line, we calculate the accessibility score by checking how many motif instances are found in the open chromatin fraction of the genome. After further normalization, these accessibility scores are compared to gene expression values for all genes via regression (Methods). To account for confounding, we use mixed model regression, where an additional random component is used with the same covariance structure as the gene expression matrix. To be considered a CSR candidate, motif accessibility of a TF must show strong association (low p-value) with the expression of the corresponding TF gene compared to other genes. The gene-level *CSR rank* of a TF is defined as the rank of its association p-value among the p-values for all genes.

## Results

### A linear mixed model approach to predict chromatin state regulators

Our approach rests on the assumption that the activity of a CSR is correlated with the amount of open chromatin in the vicinity of its potential binding sites. Both quantities can be estimated from genomic data: For the CSR activity we use its gene expression level as a proxy for the active protein concentration. The effect of this activity is approximated by the open chromatin fraction of the genome around its binding motif instances (Fig. 1). Specifically, we count the number of instances of a given TF’s binding motif in the open chromatin fraction of the genome to define a motif accessibility score. A naive approach would be to use standard linear regression between the motif accessibility score and the expression level of a given TF to identify CSR candidates. Yet, this method has an elevated type I error rate, as it does not account for confounding due to (substantial) cell line relatedness and batch effects. To overcome this limitation, we use here a linear mixed model (LMM) framework, where a random effect accounts for such confounding factors (which has been shown to work well in genetic association studies (Kang et al. 2008, 2010; Lippert et al. 2011)).

For a given motif, we use the linear mixed model framework to find the association p-value between its accessibility score and the measured expression of the TF gene. We then compare this p-value to the p-values calculated using the measured expression of each of the other genes. If confounding is controlled for, most association p-values should follow the uniform [0,1] distribution. Furthermore, if the TF is a CSR, its p-value should be low compared to other genes. We thus define the *CSR rank* of a TF as the rank of its association p-value among all genes (see example in Fig. 1). Low CSR ranks indicate strong association between motif accessibility and TF expression, suggesting that the TF is a CSR.

To apply our method, we used data released as part of the ENCODE project (Bernstein et al. 2012). Specifically, we used DHS-data as well as mRNA expression data across 109 cell lines. To calculate motif accessibility scores we used 325 TF binding motifs from the HOCOMOCO database (Kulakovskiy et al. 2013). As expected, we observed severe confounding when using standard linear regression, which was controlled using our method based on mixed model regression (Fig. 2).

**Figure 2:**
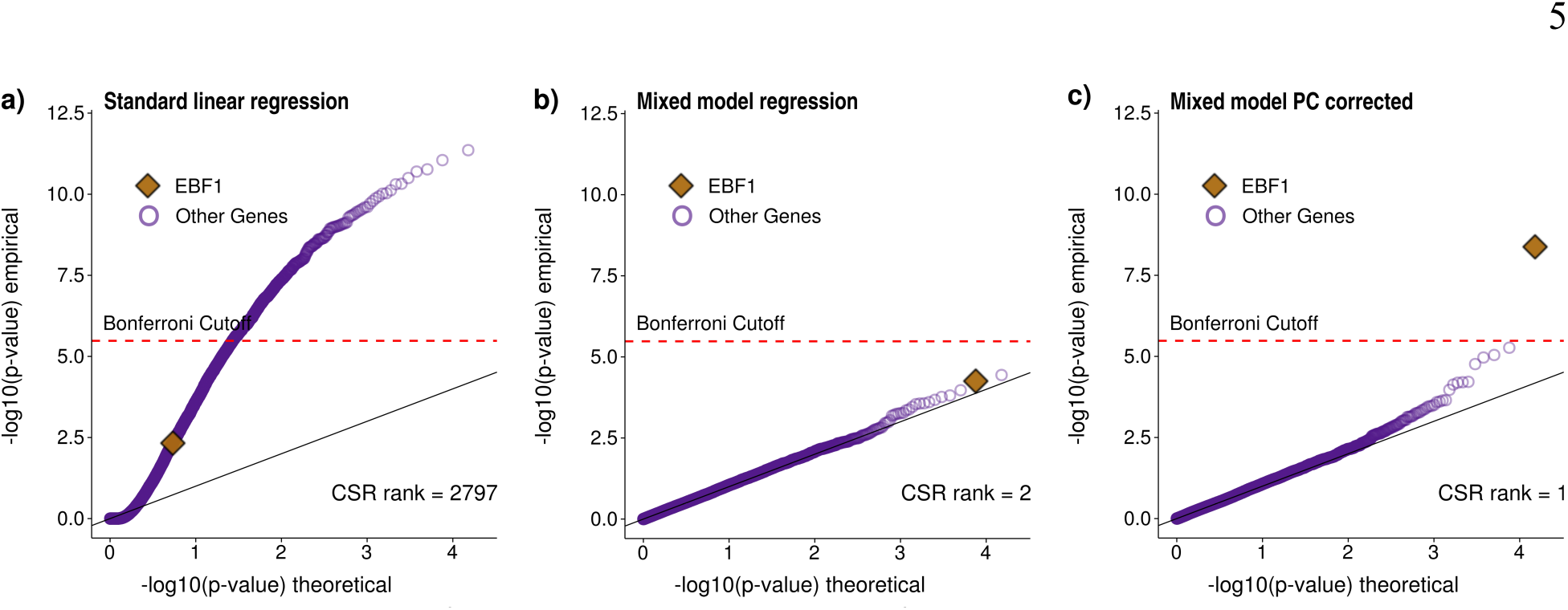
Association between motif accessibility and mRNA expression for the putative chromatin state regulator *EBF1*. Three different regression models (a-c) were used to compute association p-values between the accessibility of a given TF motif (here *EBF1*) and mRNA expression for each of the assayed 15K protein-coding genes. Results are visualized as qq-plots showing the ‒log10 transformed p-values. (**a**) Association p-values obtained using standard linear regression. Due to confounding, p-values are strongly inflated and *EBF1* motif accessibility does not show strong association with *EBF1* expression compared to other genes. (**b**) The linear mixed model (LMM) successfully corrects for confounding, with most p-values following the null distribution as expected. The association between *EBF1* motif accessibility and *EBF1* expression now ranks second among all genes and first among all TFs, although it does not pass the Bonferroni significance threshold. (**c**) Additionally controlling for the first principal component of the motif accessibility matrix corrects for a strong batch effect (Methods), which further improves the signal. Using this approach, *EBF1* motif accessibility showed the strongest association precisely with *EBF1* expression (i.e., the gene-level CSR rank equals one), suggesting that *EBF1* may be a CSR, in agreement with the literature (Treiber et al. 2010).

### ChIP-seq shows widespread binding of homologous TFs to each others motifs

Our method relies on TF motif accessibility and expression data to predict CSRs. However, evolutionarily related TFs have similar binding motifs (Jolma et al. 2010). Motif accessibility may therefore associate not only with the expression of the annotated TF, but also with the expression of a homologous TF with a similar motif. Therefore, we mapped TFs into subfamilies using the homology-based clustering *TFClass (Wingender et al. 2013)*. The 1,557 TFs were grouped into 397 subfamilies. Using a collection of 329 ChIP-seq datasets from ENCODE, we saw strong enrichment of TF motifs in ChIP-seq peaks of the TF as well as its subfamily members (Fig. 3). We therefore consider any strong association between a motif and a member of the subfamily of its TF as a signal for a CSR.

**Figure 3:**
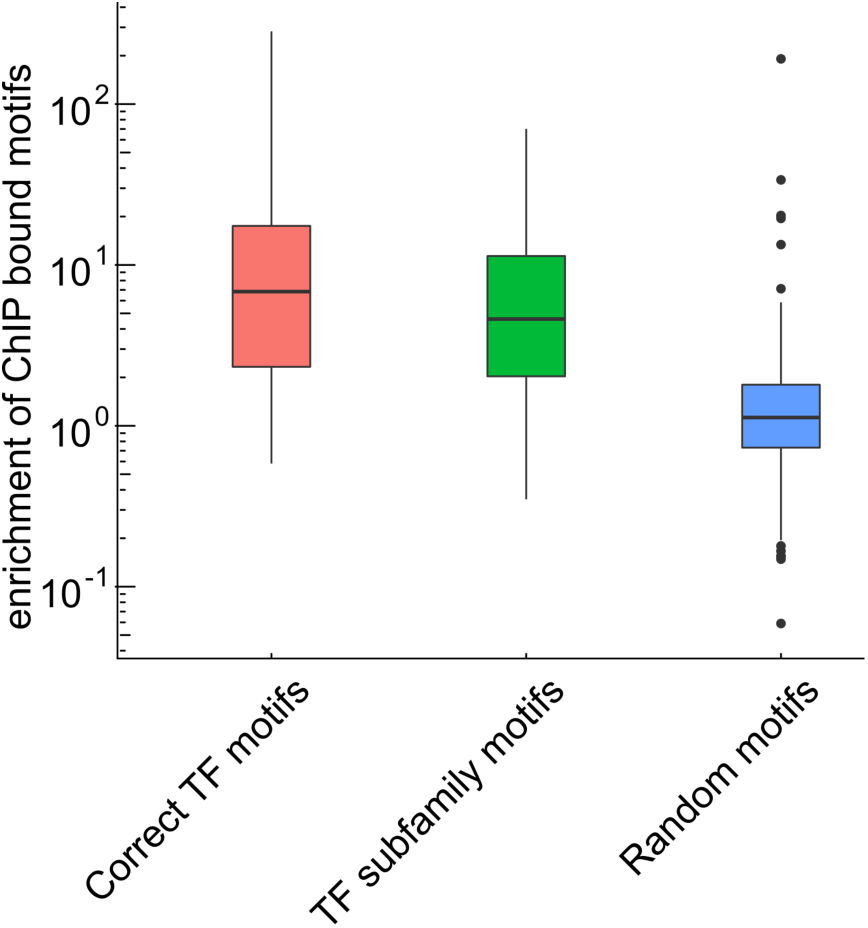
Enrichment of bound motifs for a given TF and its subfamily members. All ChIP-seq experiments from the Myers-lab released as part of the ENCODE project were downloaded and mapped to a HOCOMOCO motif (Gerstein et al. 2012). For a given ChIP-seq experiment we looked at the processed DHS-peaks in the same cell line. We partitioned DHS-peaks into two groups depending on whether they were bound by the TF (overlap with a ChIP-seq peak) or not. We then calculated both the fraction of bound and unbound DHS peaks containing a given motif. The enrichment of bound motifs was defined as the ratio of these two fractions. Results are shown from left to right for: the motifs of the TFs that were assayed in the corresponding ChIP-seq experiments (Correct TF motifs), motifs of other TFs from the same subfamily (TF subfamily motifs), and randomly sampled motifs (Random motifs). During sampling, each motif was sampled as often as the number of ChiP-seq experiment available for that motif. We see strong enrichment of TF motifs in ChIP-seq peaks of the TF as well as its subfamily members.

### Comprehensive prediction of chromatin state regulators

Next, we used the linear mixed model strategy to predict CSRs among TFs. We used 325 motifs from HOCOMOCO (after filtering motifs showing low overlap with DHS signal, see Methods). For each motif, we used the mixed model to compute its association with mRNA expression for 1,188 known TFs. Due to the redundancy of motifs within the same TF subfamily (see preceding section), we also computed *CSR ranks* at the level of TF subfamilies. To this end, we retained the most significant association p-value within each subfamily (corrected for subfamily size, see Methods and Supplementary Fig. 1). Under the null model (when TFs are not CSRs), CSR ranks should be approximately uniformly distributed across all subfamilies, so that deviation from uniformity indicates presence of CSRs.

We found strong enrichment of low CSR ranks at the subfamily level (Fig. 4, Supplementary Table 1). The enrichment was stronger when using mixed modeling instead of standard linear regression, underlining again the importance of proper control for confounding factors. When looking at the threshold that leads to 10-fold enrichment of low CSR ranks compared to uniformity (i.e., 10% false discovery rate), we found that 25% of all subfamilies have a CSR rank that falls below that threshold. These results show that many TFs do have an impact on the open chromatin fraction and can be defined as CSRs.

**Figure 4:**
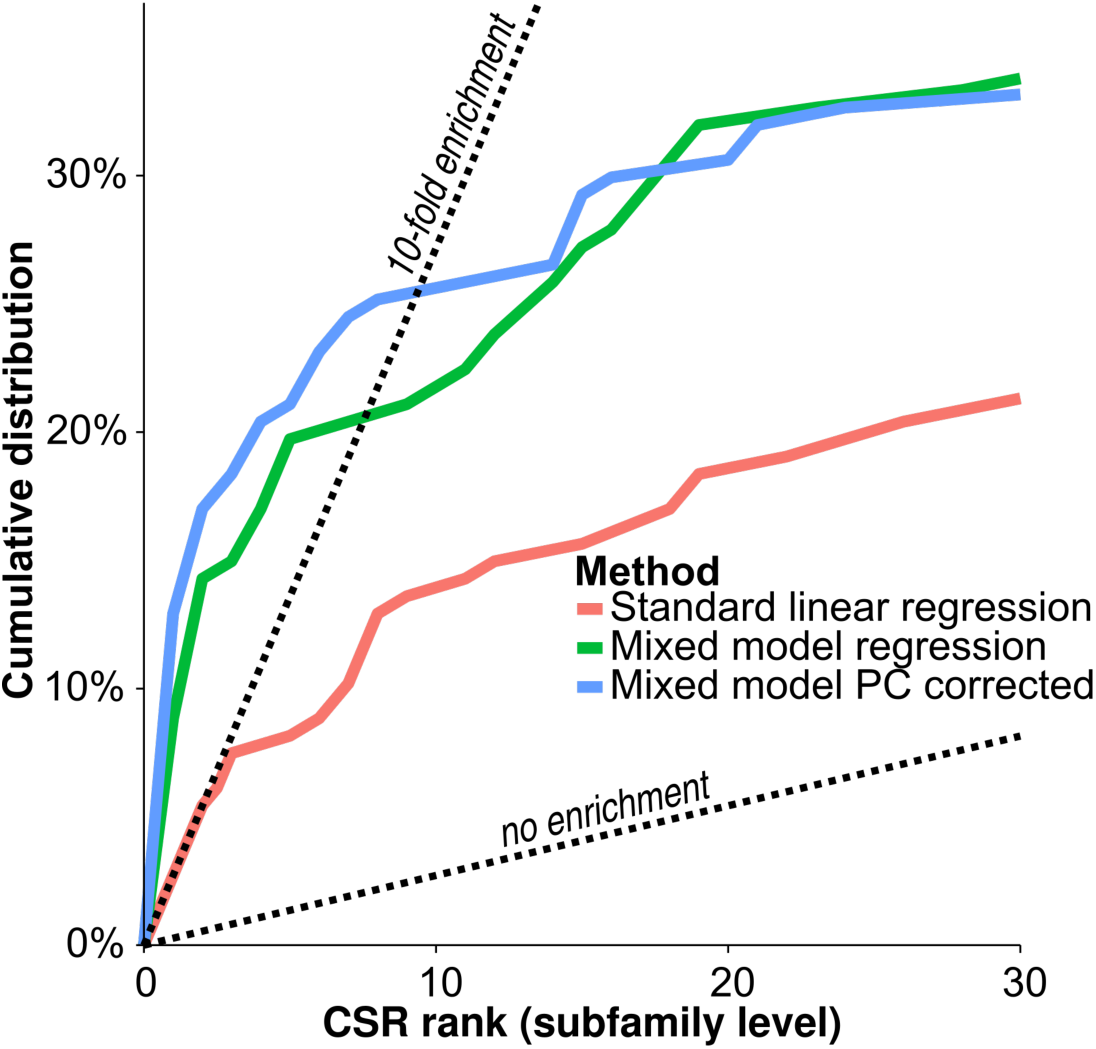
Method comparison across all subfamilies. Cumulative distribution of *CSR* ranks at the subfamily level for the 147 tested subfamilies using the three different modeling strategies: ‘standard linear regression’, ‘mixed model regression’ and ‘mixed model PC corrected’ (see legend of Fig. 2 and Methods). We see strong enrichment of low ranks implying deviation from the null hypothesis. The linear mixed modeling increases enrichment of low CSR ranks.

### Pioneer TF subfamilies enrich in predicted chromatin state regulators

As mentioned above, one well-defined class of CSRs are pioneer TFs that can bind and open closed chromatin. Therefore, subfamilies annotated to known pioneer TFs should have low CSR ranks. To test enrichment formally, we used a recently published list of established pioneer TF subfamilies (see Table 1 in Iwafuchi-Doi al. (Iwafuchi-Doi and Zaret 2014)). We asked whether these subfamilies were predicted as CSRs using our methodology. For eight subfamilies in the list for which we had the motif, six were in the fraction of ten-fold enrichment (i.e. having a CSR rank at the subfamily level below ten) (Fig. 5). To assess significance, we used permutation tests: For each of the 50,000 permutation samples, we picked eight CSR ranks from the set of subfamilies not annotated as pioneers. We summed the resulting eight CSR ranks and compared this sum to the sum of CSR ranks for the pioneer subfamilies. Using this strategy we derived a p-value of 0.0087 for the test of low CSR ranks among pioneer subfamilies. When we used a hypergeometric test at the 10-fold enrichment cutoff (Fig. 4), the p-value was even lower (0.0016). Because our approach to uncover CSRs is biased towards TFs with large mRNA expression variability (Supplementary Fig. 3), we sought to control for potential confounding introduced by the fact that the tested pioneer factors might also have large expression variability. Controlling for expression variability only slightly increased the p-values from 0.0087 to 0.024 and from 0.0016 to 0.0027, respectively (see Methods for details).

**Figure 5:**
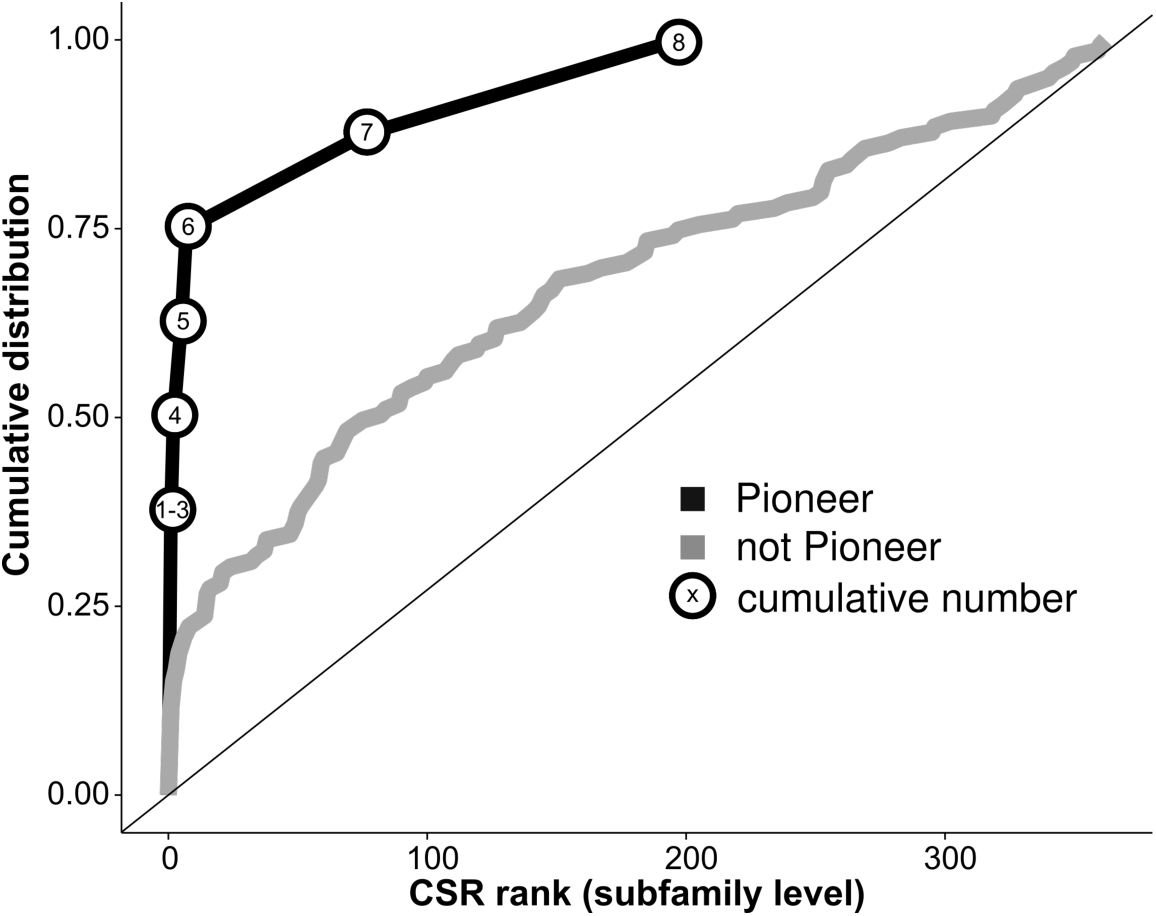
Known pioneer TF subfamilies strongly enrich in predicted chromatin state regulators. Shown in gray is a scaled cumulative distribution plot for subfamily level CSR ranks of subfamilies not annotated as pioneers in Iwafuchi-Doi (Iwafuchi-Doi and Zaret 2014). In black, we see the cumulative number of pioneer subfamilies that reached at least a given CSR rank. 6 out of 8 subfamilies show a low CSR rank which more than 3 times as many as one would expect on average when sampling from non-pioneer subfamilies.

### Downstream genes can show strong associations for activating chromatin state regulators

It is known that the activity of some TFs is mainly regulated by the level of their cofactors rather than their own protein concentration (Evans and Mangelsdorf 2014). These TFs are often present in their inactive form in the cell, which can then be quickly activated upon binding of the cofactor. This allows the cell to rapidly respond to environmental cues. An example of this phenomenon are steroid receptor TFs, which initiate transcriptional changes upon steroid hormone binding (Lu et al. 2006). In such cases, one would not expect a strong association between the mRNA expression level of a receptor TF and its motif accessibility because mRNA expression would rather be correlated to the amounts of inactive TF protein in the cell, while TF activity should depend on the strength of the environmental stimulus. However, if the TF strongly activates mRNA expression of other genes, it might be possible to determine whether the TF is a chromatin state regulator by looking at associations between the motif accessibility of the TF and the expression of its downstream genes.

To explore this strategy, we looked at associations across all genes and motifs that were below the overall Bonferroni threshold (9.6 x 10^-9^). For five out of 13 such motifs, members of the corresponding subfamily had top scores. In four further cases a gene from a TF subfamily was ranked close to the top that was highly related (i.e. part of the same family (Wingender et al. 2013)) to the motifs’ corresponding subfamily but not identical with it. This suggests that the TF subfamily clustering was too fine-grained in these cases. Surprisingly, for one motif, the significant association had a negative effect size (the negative association was observed between *NUDT11* and the motif for *RARG*). The remaining 3 motifs were all annotated to the GR-like receptors, which encompass 4 TFs (AR, *NR3C1, NR3C2, PGR*). The accessibilities of these three motifs all associated strongly with the expression of three genes (*FKBP5, ZBTB1, TSC22D3*). When using the *STRING* database to check for functional links between these genes, all genes had links to a GR-like receptor (Fig. 6) (Szklarczyk et al. 2015). In fact, all three genes are known to be glucocorticoid response genes. These results suggest that *NR3C1* might act as a CSR. For strongly activating factors, the power of the analysis can therefore be strengthened by incorporating results from downstream genes.

**Figure 6:**
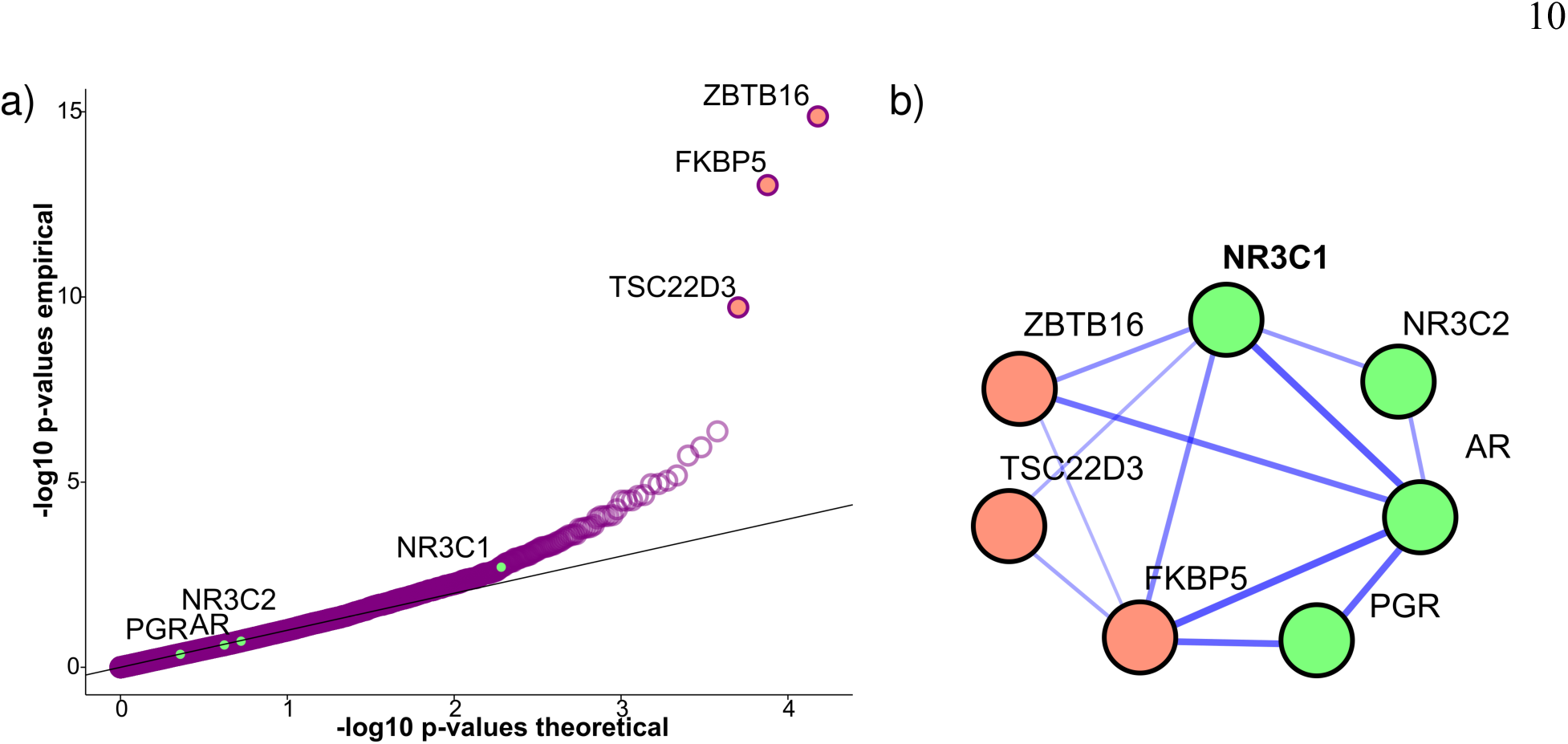
Strong associations between GR-like receptor motif and glucocorticoid response genes. a) Association results for motif accessibility of the TF *NR3C1*, which belongs to the GR-like receptor subfamily, and mRNA expression across all genes. ‒Log10 transformed p-values are shown in a qq-plot. *NR3C1* motif accessibility shows strong association with mRNA expression of three glococorticoid response genes (orange), but only weak association with expression of *NR3C1* and other GR-like receptor TFs (green). In this example, motif accessibility is strongly associated with downstream gene expression, but only weakly with expression of the TF itself. b) The network shows functional relationships among the GR-like receptor TFs (green) and the three most strongly associated genes (orange), which are all glococorticoid response genes. The strength of links shows confidence in functional relationship given in the *STRING* database. We see numerous links between the downstream glococorticoid response genes and the GR-like receptor TFs in the STRING database, confirming their functional relatedness. where *NR3C1* has the most links to associated genes.

## Discussion

It is well known that TF binding correlates with open chromatin (Thurman et al. 2012). However, for many TFs, it is not clear whether their binding is cause or consequence of open chromatin. Here, we used datasets provided by ENCODE to predict chromatin state regulator candidates, i.e., TFs that are able to establish or maintain open chromatin configurations. We devised an approach using linear mixed models to deal with the extensive confounding that one encounters in genome-wide data from heterogeneous sources. Our method uncovers a set of TFs whose expression is associated with their motif accessibility, suggesting a role in maintenance of an open chromatin configuration.

Because pioneer TFs are by definition CSRs, our predictions should include known pioneer TFs. We tested this formally for a list of pioneer TF subfamilies recently published in Iwafuchi-Doi et al^3^. Six out of eight pioneer subfamilies were indeed predicted to be CSRs: *FOXA1, GATA6, KLF4*, SOX2, *SPI1* and *TP63* were the pioneer TFs driving these signals. The two subfamilies not predicted to be CSRs were *POU5* and *CLOCK. SOX2* was the gene most strongly associated with POU5F1 motif accessibility with a low p-value of 5 x 10^-6^ (Supplementary Fig. 2). *POU5F1* acts together with *SOX2* to maintain undifferentiated states (Buganim et al. 2013). The two TFs also physically interact and a recent study proposed a model where *POU5F1* is guided by *SOX2* to target sites (Chen et al. 2014). The *CLOCK* subfamily members have a role in the cell cycle, acting as TFs in the circadian pacemakers (Fu and Lee 2003). It is possible that average mRNA expression of these TFs in unsynchronized cell lines is not a meaningful measure for their activity. In addition to the factors seen in the list, we found further factors discussed in the pioneer TF literature such as *TFAP2C, EBF1, CEBPD/B, OTX2, NFKB* and *STAT5 (Tan et al. 2011; Treiber et al. 2010; Plachetka et al. 2008; Buecker et al. 2014; Hayden and Ghosh 2012; Hagan et al. 2013*).

One limitation of our approach is that it cannot discern between open chromatin establishing TFs and open chromatin maintaining TFs. A way to discern the relative roles could be to perform overexpression and knock-down experiments followed by an open chromatin assay for the TFs found by our approach. However, this is out of the scope for the current study.

It is also clear that multiple conditions have to be met for the approach to work. First and foremost, mRNA expression has to be correlated sufficiently with protein concentration of the CSR. Typically, only a fraction of the variation in protein concentration can be explained by variation in mRNA abundances (Maier et al. 2009). Nevertheless, better power of our approach can always be achieved by increasing sample size, as long as there is at least some correlation. Further, it is reasonable to assume that our approach will perform better on TFs with a large dynamic range across cell types. This seems indeed to be the case, since most TFs predicted to be CSRs tend to have large mRNA expression variance (Supplementary Fig. 3). Sampling more and diverse cell lines could address this issue, because it should increase the dynamic range.

For some TFs, activity mainly depends on cofactors. For example, for steroid hormone receptors, hormone molecules activate a pool of inactive TF already present in the cell. In such cases measuring TF activity with gene expression measures can be misleading and one would not expect an association between the expression of a TF and the accessibility of its motif. For example, for the accessibility score of NR3C1, we saw much stronger associations with the expression levels of a small set of glucocorticoid response genes (*ZBTB16, FKBP5, TSC22D3) (Wasim et al. 2010; Galigniana et al. 2012; Rog-Zielinska et al. 2015)* than that of NRC1 itself. This difference in signal strength is in line with the activity of *NR3C1* being mainly regulated by glucocorticoid binding and not *NR3C1* gene expression levels. Interestingly, *NR3C1* was nevertheless reported to have pioneer activity (Zaret and Carroll 2011).

In summary, we exploited the rich data source of ENCODE to find TFs whose mRNA expression levels are directly linked to the open chromatin fraction of the genome. Although our approach in its current form is able to find TFs with strong associations it is also clear that increasing power by adding more cell lines would find more TFs with an association. From the current data, we would estimate that at least 25% of TF subfamilies show a low CSR rank at the subfamily level, suggesting that the regulation of chromatin state is a pervasive phenomenon amongst TFs.

## Methods

### Motif accessibility score creation

Annotated open chromatin (FDR <0.01) peaks were downloaded from the EBI website (see URL section) and trimmed to the top 90,000 peaks for each cell line. 426 motifs were downloaded from the HOCOMOCO website and aligned to the reference genome with FIMO (Grant et al. 2011). Motif occurrences with a p-value below 10^-5^ were kept for processing. For each motif, we counted the number of DHS-peaks overlapping a motif instance in a given cell line using bedops (Neph et al. 2012). Results were filtered to motifs that were present in at least 150 DHS-peaks on average, leaving 344 motifs. For a given motif, we quantile-normalized the values to follow a normal distribution yielding the raw motif-activity matrix with rows corresponding to motifs and columns corresponding to cell lines. The resulting matrix was iteratively scaled to zero mean and unit standard deviation, first row-wise (across cell lines) then column-wise, until convergence (Efron 2013; Olshen and Rajaratnam 2010). Next, we saw that the cell-line wise covariance matrix had a very large first eigenvalue, with a corresponding eigenvector that did not track well the different tissue origins of the various cell lines. Assuming that this leading principal component largely captured batch effects, we chose to regress out the first eigenvector from each row of the matrix, leading to better agreement between expression and motif accessibility correlation matrices (Supplementary Fig. 4). After this step, we quantile-normalized the data per motif to follow a normal distribution to ensure modeling assumptions of the statistical model we employed for regression were met.

To map motifs to TFs and TF subfamilies, we used the *TfClass* hierarchy (Wingender et al. 2013). Of the 344 tested motifs, we mapped 330 to a TF and its subfamily. Of these, 325 had expression data available for a subfamily member.

### Expression matrix creation

We downloaded raw expression microarray data from the GEO repository (GSE1909 and GSE15805). We background corrected and normalized using the RMA-algorithm implemented in the oligo package to process all arrays for which DHS data was also available (Irizarry et al. 2003; Carvalho and Irizarry 2010). Only the core set data was used. The data were summarized to gene level (Wells et al. 2013). Only results that had a one-to-one mapping between genes and gene probesets were kept. 15,119 genes could be annotated in this fashion. Because for many cell lines more than one experiment was conducted, we summarized multiple plates by averaging gene results across experiments. The resulting matrix was iteratively scaled to zero mean and unit standard deviation, first row-wise (across cell lines) then column-wise, until convergence (Efron 2013; Olshen and Rajaratnam 2010).

### Linear Mixed effect Model

The model proposed is

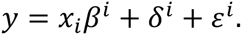

Where *y* is a vector of motif accessibility scores across *n* cell lines, *x*_*i*_ is the expression vector of gene *i*, *β*^*i*^ is the effect size of gene *i*:

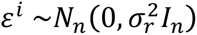

and

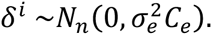

*C*_*e*_ is the covariance matrix of the *n*×*p*expression matrix:

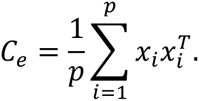

For each gene 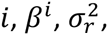 and 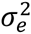 are estimated via maximum likelihood and the hypothesis *β*^*i*^ = 0 is tested via a likelihood ratio test.

### Calculating CSR ranks at the subfamily level

For each motif in HCOCOMOCO, we used the mixed model association results across all 1,188 known TFs for which we had mRNA expression data. This yielded a matrix of association p-values for all pairs of 325 motifs (belonging to 147 *TFClass* subfamilies) and 1188 TFs (belonging to 368 *TFClass* subfamilies). Due to the fact that homologous TFs have similar binding motifs, we sought to aggregate results into CSR *ranks* at the subfamily level (Supplementary Fig. 1). To achieve this, we reduced the 325x1188 motif-TF association matrix to a 147x368 association matrix of motif subfamilies and TF subfamilies. In practice, for each motif subfamily-TF subfamily pair we collected the most significant p-value among all motif-TF pairs in these subfamilies and multiplied it with the total number of such motif-TF pairs to correct for subfamily size. Finally, for each motif *subfamily*, we ranked the adjusted p-values across all TF subfamilies and defined its *CSR rank* as the rank of its corresponding TF subfamily.

### Controlling pioneer enrichment for expression variation

To control pioneer enrichment for mRNA expression variation, we first calculated the expression variance of each TF across all cell lines. The distribution of variance values was transformed to follow a standard normal distribution. We then used the maximal expression variance observed for any TF in each subfamily. To assess significance, we used permutation tests: we sampled 8 non-pioneer subfamily level CSR ranks 50,000 times. However, subfamilies were not sampled uniformly: We sampled 4 nonpioneer subfamilies with maximal expression variance between the 0th and the 50th quantile of the 8 pioneer subfamilies, and 4 non-pioneer subfamilies with maximal expression variance between the 50th and the 100th quantile of the 8 pioneer subfamilies.

## URLs

Code for reproduction (including scripts for data download) is available at: https://github.com/dlampart/csrproject

DHS peaks were downloaded from: http://ftp.ebi.ac.uk/pub/databases/ensembl/encode/integration_data_jan2011/byDataType/openchrom/jan2011/fdrPeaks/

## Acknowledgements

ZK received financial support from the Leenaards Foundation, the Swiss Institute of Bioinformatics and the Swiss National Science Foundation (31003A-143914) and SystemsX.ch (51RTP0_151019). SB received funding from the Swiss Institute of Bioinformatics, the Swiss National Science Foundation (grant FN 310030_152724 / 1) and SystemsX.ch through the SysGenetiX project.

